# The impact of preprint servers in the formation of novel ideas

**DOI:** 10.1101/2020.10.08.330696

**Authors:** Swarup Satish, Zonghai Yao, Andrew Drozdov, Boris Veytsman

## Abstract

We study whether novel ideas in biomedical literature appear first in preprints or traditional journals. We develop a Bayesian method to estimate the time of appearance for a phrase in the literature, and apply it to a number of phrases, both automatically extracted and suggested by experts. We see that presently most phrases appear first in the traditional journals, but there is a number of phrases with the first appearance on preprint servers. A comparison of the general composition of texts from bioRxiv and traditional journals shows a growing trend of bioRxiv being predictive of traditional journals. We discuss the application of the method for related problems.

## 1 Introduction

A paper submitted to a journal goes through several stages: peer review, editorial work, copyediting, publication. This leads to a long waiting time between submission and the final publication (Powell, 2016). The situation is especially bad in the life sciences, where the waiting time between submission and publication approaches the duration of a traditional PhD study, creating serious difficulties for young scientists (Vale, 2015). While this is frustrating for scientists whose recognition and promotion often depend on the publication record, it is also bad for science itself, significantly slowing down its progress (Qunaj et al., 2018).

Preprint servers were offered as a means of accelerating science (Berg et al., 2016; Desjardins-Proulx et al., 2013; Sarabipour et al., 2019; Schloss, 2017; Lauer et al., 2015; Peiperl, 2018), especially in the wake of COVID-19 epidemics (Krumholz et al., 2020). The discussion about the benefits (and dangers) of preprint is no longer confined to the scientific literature, coming to the pages of popular newspapers (Eisen and Tibshirani, 2020).

The benefits of preprints for accelerating science are often raised in the discussions between regulatory agencies, funders, scientists and publishers. Thus a method to objectively assess them is important. One way to do this assessment is to look at a new important idea and to measure whether it first appears in a traditional journal or on a preprint server.

An implementation of this approach requires one to define what is an important idea, and how to find the time of appearance for it. This is the goal of our work.

The definition of novelty in science and the methods to determine and predict novelty have a long history discussed in the next section. In this work we use a very simple approach (Garfield, 1967; Latour and Woolgar, 1986): new ideas correspond to new terms. Thus if we find new words and phrases in scientific papers, we can surmise the appearance of new ideas.

The definition of the time of appearance for a new idea is not trivial. It is not enough to register the first mention of a term. First, some hits might be erroneous, and give us false positives. On the other hand, we might miss some mentions of a term due to the incompleteness of the corpus. Therefore a more subtle method to determine the time of appearance is needed. In this work we offer a Bayesian approach to this problem.

Based on our definitions of novelty and the time of appearance for novel ideas we compare the time of appearance for several novel ideas in the papers published in bioRxiv/medRxiv https://www.biorxiv.org/ and PubMed Central full text collection https://www.ncbi.nlm.nih.gov/pmc/.

## 2 Related Works

Novel ideas and breakthroughs are among the central concepts for the science of science. A number of studies propose different ideas to quantify originality in science and technology (Cozzens et al., 2010; Alexander et al., 2013; Rzhetsky et al., 2015; Rotolo et al., 2015; Wang et al., 2016; Wang and Chai, 2018; Shibayama and Wang, 2020) or their impact on the other works (Shi et al., 2010; Shahaf et al., 2012; Sinatra et al., 2016; Hutchins et al., 2016; Wesley-Smith et al., 2016; Herrmannova et al., 2018b,a; Zhao et al., 2019; Bornmann et al., 2019; Small et al., 2019). The prediction of breakthroughs, scientific impact and citation counts is a well developed area (Schubert and Schubert, 1997; Garfield et al., 2002; Dietz et al., 2007; Lokker et al., 2008; Shi et al., 2010; Uzzi et al., 2013; Alexander, 2013; Klimek et al., 2016; Tahamtan et al., 2016; McKeown et al., 2016; Clauset et al., 2017; Peoples et al., 2017; Salatino et al., 2018; Dong et al., 2018; Iacopini et al., 2018; Feldman et al., 2018; van den Besselaar and Sandström, 2018; Klavans et al., 2020). However, the question asked in these works is different from the one we ask. Most of the researchers tried to determine what makes a work original or impactful, and how to predict originality or impact. Our question is the following: suppose we know a certain idea is novel (or impactful). Can we pinpoint a moment in time when this idea appeared, and where did it appear?

Sentence level novelty detection was a topic of novelty tracks of Text Retrieval Conferences (TREC) from 2002 to 2004 (Soboroff and Harman, 2003; Harman, 2002; Clarke et al., 2004; Soboroff and Harman, 2005). The goal of these tracks was to highlight the relevant sentences that contain novel information, given a topic and an ordered list of relevant documents. At the document level, Karkali et al. (2013) computed novelty score based on the inverse document frequ function. Another work by Verheij et al. (2012) presents a comparison study of diffe detection methods evaluated on news a language model based methods perform better than the cosine similarity based ones. Dasg (2016) conducted experiments with information entropy measure to calculate novelty of a document. Again, the work that we present here significantly differs from the existing novelty detection methods since we use the novelty detection as a starting point rather than a goal. We first get candidate phrases from the documents with the most appropriate key phrases extraction method, and then determine the appearance timing of these phrases.

Another field that is relevant to our research is the detection of change points in a stream of events. Change point detection (or CPD) detects abrupt shifts in time series trends that can be easily identified via the human eye, but are harder to pinpoint using traditional statistical approaches. The research in CPD is applicable across an array of industries, including finance, manufacturing quality control, energy, medical diagnostics, and human activity analysis. There are many representative methods of CPD. Binary segmentation (Bai, 1997) is a sequential approach: first, one change point is detected in the complete input signal, then series is split around this change point, then the operation is repeated on the two resulting sub-signals. As opposite to binary segmentation, which is a greedy procedure, bottom-up segmentation (Fryzlewicz, 2007) is generous: it starts with many change points and successively deletes the less significant ones. First, the signal is divided in many sub-signals along a regular grid. Then contiguous segments are successively merged according to a measure of how similar they are. Because the enumeration of all possible partitions is impossible, Pelt (Killick et al., 2012) relies on a pruning rule. Many indexes are discarded, greatly reducing the computational cost while retaining the ability to find the optimal segmentation. Window-based change point detection (Aminikhanghahi and Cook, 2016) uses two windows which slide along the data stream. Dynamic programming was also used for this task (Truong et al., 2020). In this work we propose a simple Bayesian approach to the detection of change points, which seems to give intuitively reasonable results for our purpose.

## 3 Bayesian Approach to Novelty Detection

Our proposed Bayesian Approach to Novelty Detection (BAND) finds the time interval *τ* that maximizes the observed series of publication frequency for a phrase (Figure 1). Paper publications are events, so it is reasonable to assume that the number of publications *n* in a unit interval at the time *t* containing the given phrase *z* follows a Poisson distribution. The joint density function for publication frequency is given by the equation

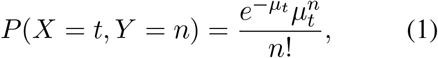

where *μ*_*t*_ is modeled by a piecewise linear function of *t*:

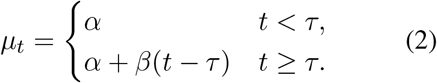

**Figure 1:**
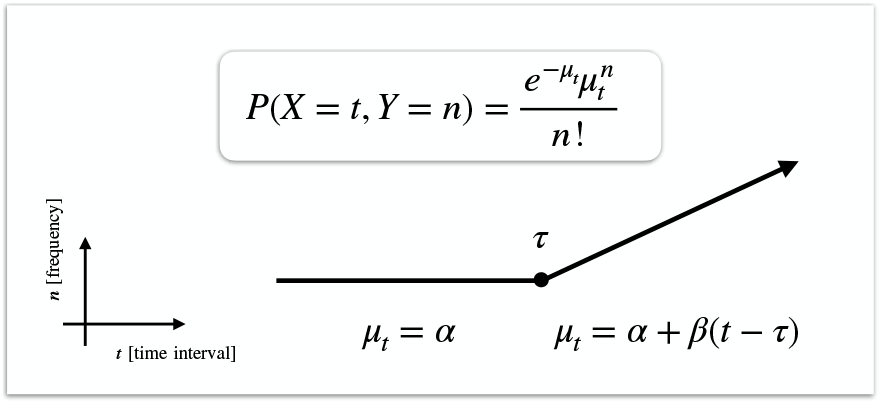
Our Bayesian Approach to Novelty Detection (BAND). The goal is to find the inflection point *τ* that indicates the earliest point attributed to the rapid research growth associated with a novel idea.

Prior to *τ*, we expect the number of publications containing the given phrase to be small, ideally zero. The parameter *α* > 0 controls for the noise (misattributed papers, improper or ambiguous usage of the phrase, etc.). After the moment *τ*, we expect the steady grow of phrase popularity with the rate *β*. In other words, *τ* is the point in time ehen the phrase begins to be adopted.

We consider each phrase independently and use Bayesian modeling to find the most probable parameters *α*, *β*, and *τ* given the observed data using the standard Bayesian approach

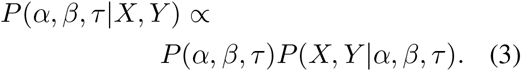

We use a flat uniformative prior with

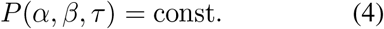

Our motivation for finding *τ* using BAND is to compare the impact of different publication venues (i.e. preprint servers and peer-reviewed journals) that cover overlapping research topics. For a single phrase *z*, we use the observed data from two sources and estimate the posterior probability *P* (*α, β, τ*) for each source separately. Then we run a simulation to find the 95% confidence interval of *τ* using the following procedure:

1. For a single data source, compute *P* (*α, β, τ*) for all possible configurations on a grid.
2. Sample a large number of triplets (*α, β, τ*) from the posterior computed in Step 1.
3. Remove 2.5% of the triples with the highest value of *τ*, and 2.5% of triplets with the lowest value of *τ*. The probability for *τ* to lie in the remaining interval can be estimated as 95%.

In order to compare two sources, we create two sets of tuples, one for each source. Then we randomly draw a tuple from the first set and a tuple from the second one. For each pair *i* we compute *δ*_*i*_, the expected difference between the two sources where 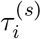 is the *i*-th sample in source *s*:

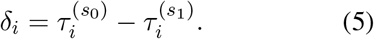

The conclusion about the priority is based on distribution of *δ* around zero. If *δ* > 0 for the majority of the pairs, then the phrase gains traction first on source *s*_0_. Otherwise the second source wins.

## 4 Experimental Setup

In this section we describe our data collection procedure and background on methods of text processing and assessing data quality. Our code for running experiments is publicly available.^1^

### 4.1 Data Sources (Publication Venues)

We consider two types of publication venues: peer-reviewed journals and preprint servers. There are now many preprint servers for various areas of science, including arXiv, bioRxiv, medRxiv, PsyArXiv, SocArXiv, ChemRxiv, AgriRXiv, and others. To compare a preprint server to a traditional venue one needs a large open access collection of traditionally published papers. In biomedical sciences there is a huge Pubmed Central Open Access Dataset described below, which drove our choice for bioRxiv/medRxiv as a comparison venue. Another reason for this choice of data sources is that one of our organizations, Chan Zuckerberg Initiative, has a special interest in biomedical sciences in general and bioRxiv & medRxiv in particular.

PubMed is a central repository for biomedical papers published in peer-reviewed journals. It contains over 26 million journal publications. Abstracts are publicly available for all papers, and for a subset (the PubMed Central Open Access Dataset with over 1.6 million papers) full texts are available.^2^

In contrast to PubMed, bioRxiv is a preprint server for the biological sciences. Papers published there are not required to pass a strict and lengthy review process. BioRxiv hosts over 70,000 full text articles, each open to the public. Recently medical papers were separated into a special server medRxiv. Since the search engine provided by bioRxiv can search both servers, below we use the term “bioRxiv” for the longer, but more correct term “the union of bioRxiv and medRxiv papers”.

PubMed and bioRxiv cover the two representative categories of publication venue. We also include data sources using information from COVID-19 Open Research Dataset Challenge (Wang et al., 2020).

### 4.2 Text Processing and Data Collection

In our experiments and analysis we leverage a large collection of phrases collected in an unsupervised way using TextRank (Mihalcea and Tarau, 2004; Nathan, 2016). The procedure to extract the phrase is the following:

1. First we extract all candidate phrases from bioRxiv abstracts using TextRank. This results in 1,587,408 phrases.
2. We filter the phrases from Step 1 to the 239,608 phrases by eliminating phrases that were detected by TextRank only once.
3. For each phrase from Step 2 we generate monthly time series data for both PMC and bioRxiv using full text.^3^

We use this data collection procedure and recommendations from CZI biomedical curators to create three groups of phrases:

a. Common phrases: 20 banal phrases manually selected that are also extracted by TextRank (includes ‘medical history’, ‘heart disease’, ‘x-ray’, etc.).
b. Novel phrases: 20 phrases selected by experts (includes ‘mass cytometry’, ‘gene editing’, ‘fluorescence activated cell sorting’, etc.).
c. Top extracted phrases: The top 7000 phrases extracted by TextRank determined by average importance score.

All common and novel phrases, and a subset of the top ranking phrases are shown in Appendices A.2, A.1.2, and A.1.1.

### 4.3 Baselines for finding τ

To assess the effectiveness of BAND for finding *τ* we include a strong baseline in our experiments. In our analysis (Section 5.2), we compare these methods to BAND not only for finding t he first clear inflection point, but also how relevant *τ* is for the end goal of novelty detection and comparing research impact.

The baseline we include is Window-based Change Point Detection (see Section 2). It works by maximizing the discrepancy measuring function

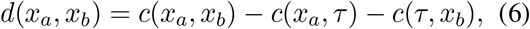

which is large when the left segment *c*(*x*_*a*_, *τ*) is dissimilar from the right segment *c*(*τ*, *x*_*b*_).

Window-based Changepoint Detection (WCPD) is flexible and has been successful when applied to many tasks. However, for a number of novel phrases (see the first two examples on Figure 3) it gives intuitively unsatisfactory results for the detection of the idea onset. The reason is, WCPD tries to find the point where the publication frequency significantly changes, which often corresponds to the moment the idea is widely adopted. Our task, on the other hand, is to find the point when the idea appears, which is a different problem. Therefore one may expect BAND to work better for the cause of the novelty detection because its model of growth starting from zero might be more suitable to describe the publication frequency than the generic model of WCPD.

We found necessary to run WCPD using the finite difference approximation of the publication frequency gradient to obtain reasonable predictions.

We use the implementation of WCPD provided in ruptures (Truong et al., 2020) with an *L*^2^-cost function (*c*), window size of 10, penalty of 1, and no specification on how many changepoints to return.^4^ In our figures we display all changepoints from WCPD to illustrate these changepoints do not solely identify the point before rapid growth in idea adaptation. On the other hand our method (BAND) consistently finds changepoints that match this criterion.

## 5 Results and Analysis

In this section we discuss the following hypotheses and research questions:

- Can we find the point *τ* in time when a phrase is determined novel using our Bayesian Approach to Novelty Detection (BAND)?
- Is the value of *τ* effective for comparing the impact of two publication venues?
- Do our findings verify that preprint servers are having a positive impact on the development of novel ideas?
- How effective is publication frequency for distinguishing from novel and banal phrases?

Below we answer each of these questions.

### 5.1 Are pre-print servers accelerating research?

The common wisdom is the preprint servers have positive impact on research, by making research available openly and quickly. We attempt to verify that ideas develop faster on bioRxiv rather than on PubMed. Our results indicate that this might be true in some, but not all, cases. For some phrases *δ* leans positive (see Figure 2 top), indicating the relevant phrase and presumably the novel idea appeared first on bioRxiv. However, the opposite is true more often than not (see Figure 2, bottom).

**Figure 2:**
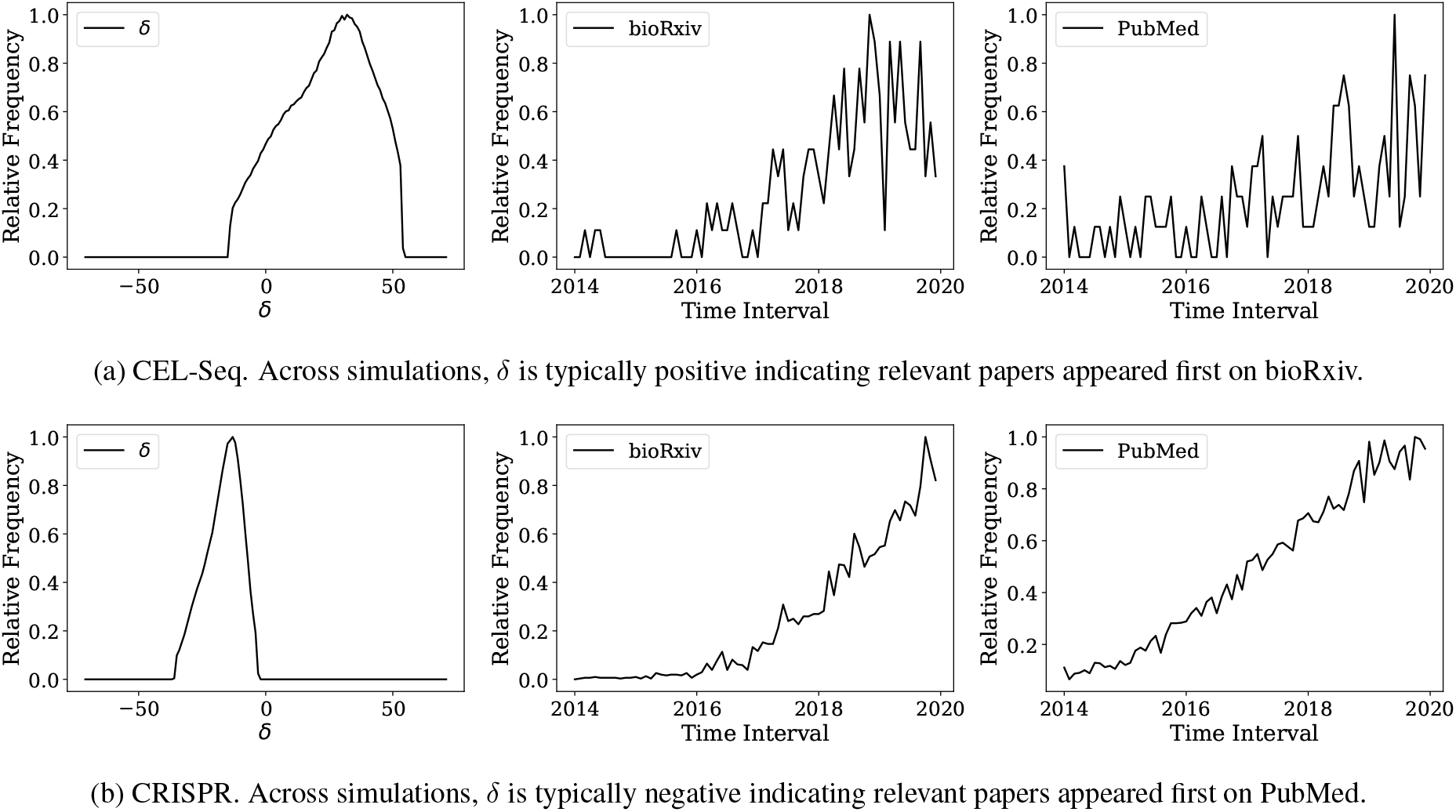
The distribution of *δ* (left column), publication frequency on bioRxiv (center), and publication frequency on PubMed (right) for two phrases

Why is PubMed frequently the first place that novel ideas appear? One reason might be that bioRxiv is relatively new, did not gain enough traction yet, and its benefits are not widely appreciated. One can even say it is surprising and encouraging that *some* novel ideas appear first on bioRxiv despite the it being a relative newcomer. Thus one interpretation of our finding is that while preprint servers are already having a positive impact on research, there is still a potential for the growth. In this case we expect that in the future novel ideas will appear first on bioRxiv at a higher rate. We further check this assumption in Section 5.3.

### 5.2 Is BAND effective at determining τ ?

We assume that curves of publication frequency of novel ideas follow a particular shape—they are relatively flat followed by a growth period. This assumption is built in the design of BAND. To verify BAND’s effectiveness, we use a set of novel phrases provided by the team of biomedical curators at Chan Zuckerberg Initiative (CZI), extract their publication frequency data from PubMed, and calculate *τ* using BAND. Qualitatively, we see in Figure 3 that BAND results agree with the intuition about the novelty onset.

**Figure 3:**
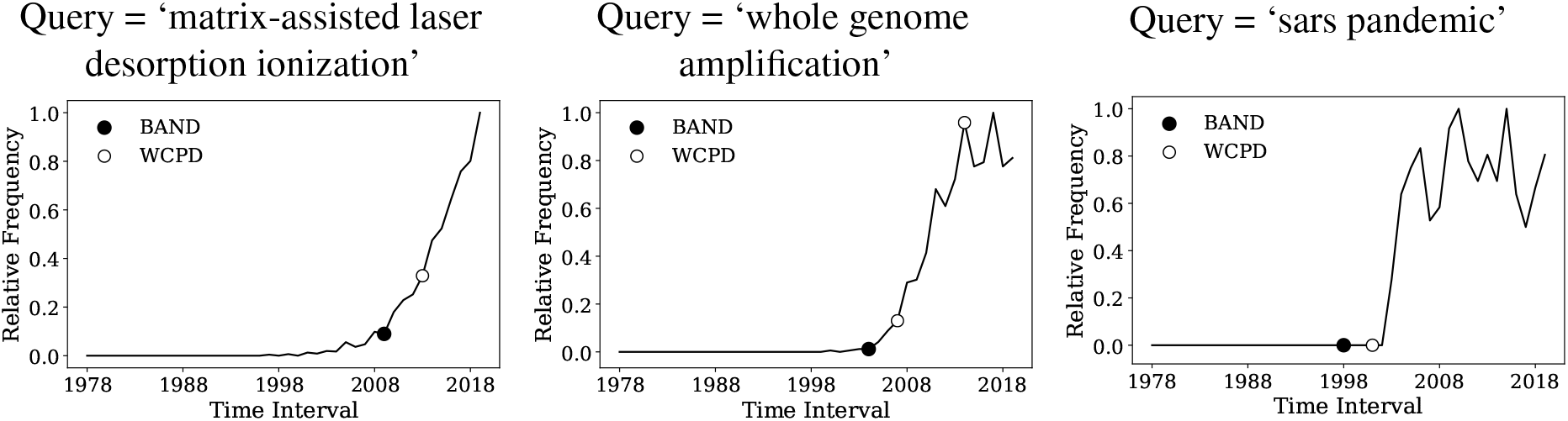
BAND often finds a more desirable point of inflection than WCPD (left). In addition, WCPD may return multiple points (center). However, in cases when the evolution of a term does not follow our model, WCPD is more effective (right).

As discussed in Section 4.3, WCPD is another technique for finding points of inflection. WCPD is a useful method because it finds inflection points without requiring any prior model of the data. As expected, if the evolution of data follows our model of growth, BAND gives a better estimate for the novelty onset (Figure 3, left and center). On the other hand, if the data do not follow BAND model, BAND is not supposed to work well, and assumption-free models like WCPD might work better. An example is shown on (Figure 3, right), where a period of growth is followed by a plateau rather than the growth assumed by BAND model.

### 5.3 Are pre-print servers mature?

A potentially confounding variable in our experimental setup is the relative recency in the establishment of bioRxiv (2013) compared to PubMed (1996). This motivates us to measure the correlation between the publication frequency of these two data sources. We proceed by looking at three groups of phrases: (a) common phrases found in medical terminology, (b) novel phrases provided by experts, and (c) highest scoring phrases according to TextRank.

First, we aggregated the last 6 years of data. We found the similarity between PubMed and bioRxiv for common phrases and phrases extracted from TextRank most similar when comparing the early snapshot of bioRxiv with the later snapshot of PubMed (Figure 4). Thus for these phrases bioRxiv is predictive of PubMed. On the other hand, phrases selected by experts (Section 5.2) tend to appear on PubMed first (Figure 4, center).

**Figure 4:**
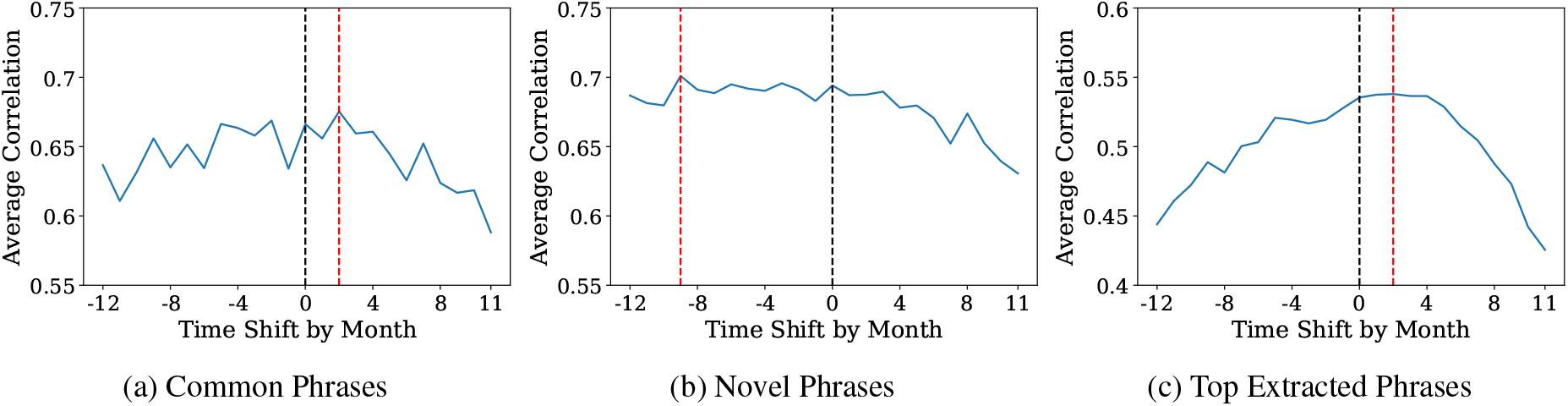
Coarse-grained correlation analysis. We report average correlation between PubMed and bioRxiv for publication frequency in a 6-year window. We shift the window for bioRxiv one month at a time, where positive values on the x-axis correspond to shifts back in time and negative values indicate shifts forward in time. If at the highest value (indicated by the dotted red line) the offset is positive bioRxiv is a predictor of PubMed’s content. If the offset is negative, the opposite is true. We perform this analysis for 3 subsets of terms: common phrases (a), novel phrases (b), and phrases from TextRank (c) starting in January 2014.

Taking into account that bioRxiv is still evolving, we performed a more fine-grained analysis where results are only aggregated across 2 years instead of 6. This also allowed us to study how the content alignment between PubMed and bioRxiv has changed over time. The result is shown on Figure 5. According to this figure, as bioRxiv matures, it becomes a better leading indicator of PubMed. Perhaps more critically, it also shows that the content alignment is indeed changing.

**Figure 5:**
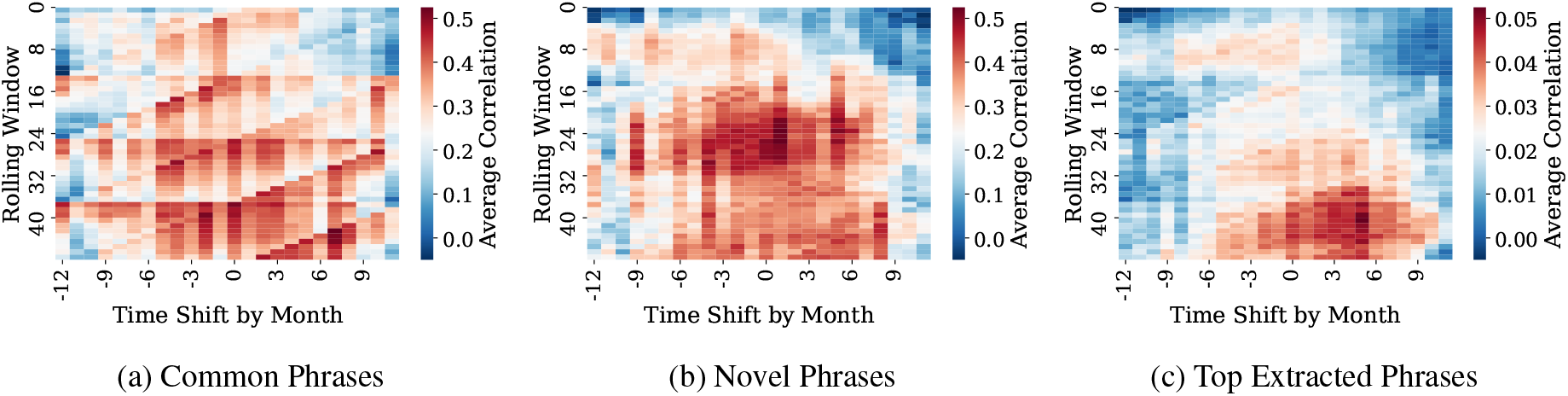
Fine-grained correlation analysis. We perform the same analysis as in Figure 4, except with a smaller window (two years) and shifting both bioRxiv (*x*-axis) and the starting month (*y*-axis). If *x* = 1 and *y* = 3, then the start month is April 2014 for PubMed and March 2014 for bioRxiv. The data are shown on a grid with darker red indicate high correlation and dark blue indicating low or inverse correlation. In general, we see higher correlation between bioRxiv and PubMed as bioRxiv matures.

The analysis we provide in this work at best is a glimpse into the relationship between pre-print servers and peer-reviewed journals. It will likely change as bioRxiv continues to mature.

### 5.4 Qualitative analysis for the spread of ideas during virus outbreaks

During natural disasters such as virus outbreaks, scientific progress towards understanding diseases and their cures is critical, warranting fast dissemination of ideas and results of research. Pre-print servers are particularly well suited to this end. Thus we compare bioRxiv to PubMed for five recent virus outbreaks, some of which took place prior to the founding of bioRxiv.

Each virus outbreak was analyzed using a composite of publication frequency of multiple related phrases. For example, values for ‘sars-cov2’ and ‘covid-19’ are aggregated in the COVID-19 plot. The five relevant outbreaks are listed below:

- Prior to bioRxiv establishment (before November 2013, Figure 6, top): MERS and SARS.
- After bioRxiv establishment (Figure 6, bottom): Zika, Ebola, and COVID-19.

**Figure 6:**
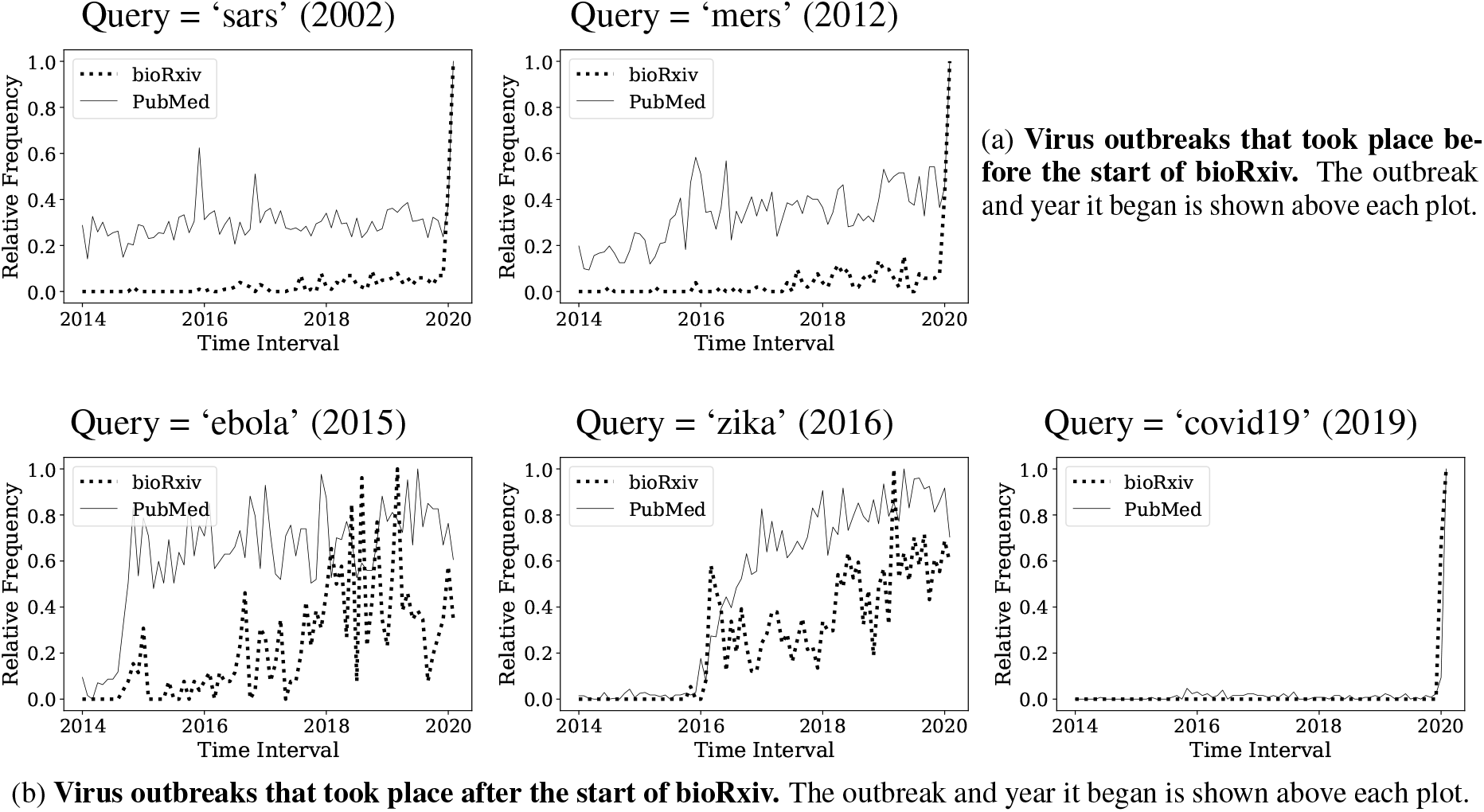
Outbreaks that occurred before bioRxiv became widespread (top row, a) with publishing activity plateaued in recent years v. outbreaks occurring after bioRxiv was founded (bottom row, b) with recent growth in research activity.

The first group of outbreaks exhibits the expected behavior: bioRxiv activity is fairly minimal given that the growth of research on those topics had begun to saturate by the time bioRxiv was formed. But the second group exhibits a different behavior. For Ebola outbreak, which took place in 2015, the activity in bioRxiv is fairly low compared to PMC. For Zika (2017) and Covid-19 (2019) bioRxiv has an early activity spike.

Through this analysis we can see that bioRxiv has become increasingly important in emergency situations during recent years.

## 6 Future Directions

Our work serves as a first attempt for discovering research impact of publication venues using full text analysis. One notable assumption we make is that all phrase mentions are treated equally. In the future, we may want to distinguish how a phrase is being used. For example, if the phrase of interest is *z*, then a research paper may write about works extending *z*, but alternatively it may simply discuss techniques similar to *z*. Similar analysis has provided useful for citations (Jurgens et al., 2018).

Furthermore, our approach leverages TextRank to find many relevant phrases, but we do not cluster phrases, so two or more phrases with similar meaning will be treated separately (i.e. FACS and Fluorescence Activated Cell Sorting). A simple alias table or string similarity extension (Tam et al., 2019) would be a clear improvement. Leveraging high precision concept extraction systems (King et al., 2020) might improve clustering even more.

Another approach would be to use a better proxy for ideas than textual phrase, like concepts or topics extracted from the corpus.

## 7 Conclusions

We introduce a Bayesian model for novelty detection (BAND), and use this model to investigate how quickly new ideas form on pre-print servers compared to peer-reviewed journals. Our findings indicate that novel phrases, which we use as a proxy for new ideas, in most cases appear on pre-print servers and in peer reviewed journals. In some cases, novel phrases appear on pre-print servers first. In many cases the content of preprints is a predictor of the content of peer reviewed journals. As the preprint servers mature, this feature becomes more prominent. When a fast review time is in high demand (such as during epidemic outbreaks), pre-print servers have a high utility, and the related novel phrases appear on pre-print servers first.

## Acknowledgments

The authors are grateful to Bill Burkholder (Chan Zuckerberg Biohub) who suggested this research, and to Barbara Vidal & Michaela Torkar (Chan Zuckerberg Initiative) for their expert help in selecting the phrases for comparison.

This research was initiated at the University of Massachusetts Amherst Industry Mentorship Program.

## A Appendices

## A.1 Summary of phrases found in data sources

In our experiments and analysis we primarily consider two data sources: bioRxiv (representative of pre-print servers) and PubMed (representative of peer-reviewed journals). We extract phrases using TextRank as described in Section 4.2. In this section of the appendix, we summarize the phrases considered in the experiments.

## A.1.1 Phrases from TextRank

TextRank extracted 1,587,408 phrases from bioRxiv abstracts. We further filter this list to 239,608 by only including phrases that were detected by TextRank more than once. This does not necessarily mean that the phrases that were filtered only occur in the bioRxiv abstracts a single time but that TextRank only found it to be a key phrase once. The distribution of phrase lengths is shown in Figure 7. A sample of phrases is listed in Table 3.

**Figure 7:**
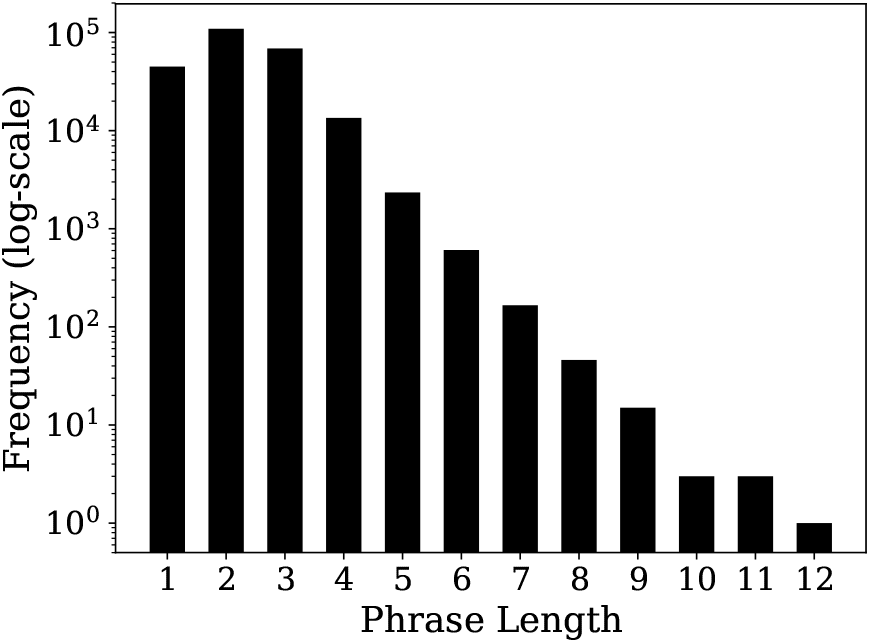
Frequency of various phrase lengths that were extracted from our corpus using TextRank.

## A.1.2 Phrases from Experts

To validate the effectiveness of BAND, we use a small set of curated phrases provided by experts at CZI. Those results are discussed in detail in Section 5.2. The complete list of phrases is shown in Table 1.

**Table 1:** The list of phrases used in Section 5.2. These phrases were provided by experts at CZI.

|  |  |
| --- | --- |
| 10X Genomics | ELISA |
| chromosome conformation | enzyme linked immunosorbent assay |
| CRISPR | Exome |
| CyTOF | FACS |
| electrospray ionization | Fluorescence Activated Cell Sorting |
| Mass Spectroscopy | Matrix-assisted laser desorption ionization |
| Proximity Extension Assay | scATAC-seq |
| Whole Genome Amplification | MALDI |
| gene editing | Mass cytometry |
| laser capture microdissection | Microfluidics |
| LINNAEUS | Translating Ribosome Affinity Purification |

## A.2 Common phrases used in bio-medicine

In conducting the correlation study, explained in Section 5.3, we used a set of phrases that can be considered to be commonly used in biomedical literature. The complete list can be found in the Table 2.

**Table 2:** The list of common phrases used in our correlation analysis in Section 5.3. These phrases are phrases that can be considered common in medical literature

|  |  |
| --- | --- |
| antibiotic | compound fracture |
| medicine | anti-inflammatory |
| physical therapy | trauma care |
| chemotherapy | medical history |
| kidney dialysis | intensive care unit |
| organ transplant | nervous system |
| X-Ray | digestive system |
| MRI | digestive tract |
| general practitioner | urinary tract |
| scar tissue | heart disease |

**Table 3:**
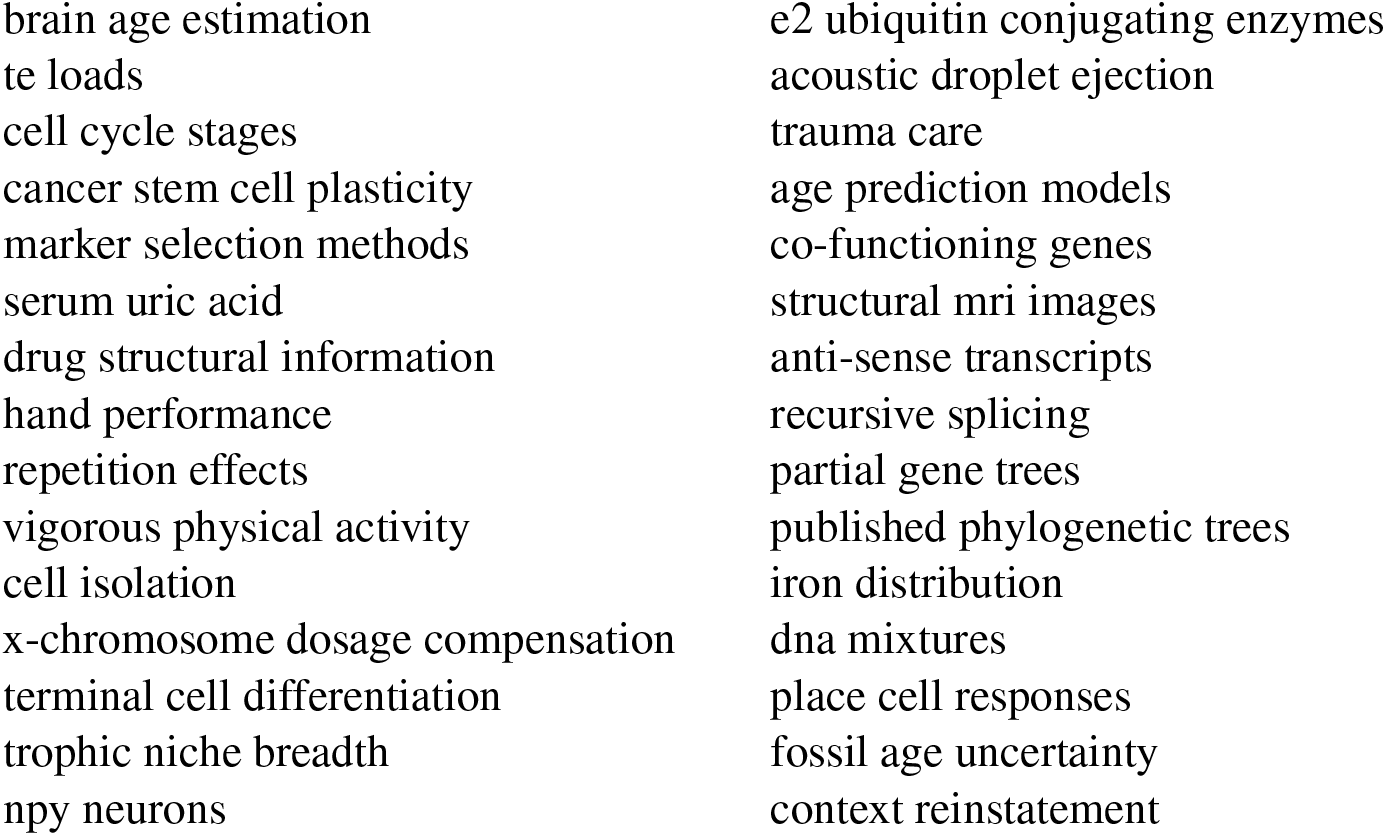
The list of top ranked phrases from TextRank used in our correlation analysis Section 5.3.

## A.2.1 Phrases for Crises

Pre-print servers are especially useful during crises such as pandemics. We chose 5 recent virus out-breaks to study their response which is discussed in Section 5.4: SARS, MERS, Ebola, Zika, and Covid-19. We made one compound time series curve for each virus’s frequency based occurrence in PubMed and bioRxiv. Table 4 shows two lists. The phrases are obtained using all ordered combinations of entries from column 2 and column 3. For example we can combine “SARS-Cov” and “epidemic” to form “SARS-Cov epidemic”. Each epidemic is associated with a set of phrases to form one compound set of phrases.

**Table 4:**
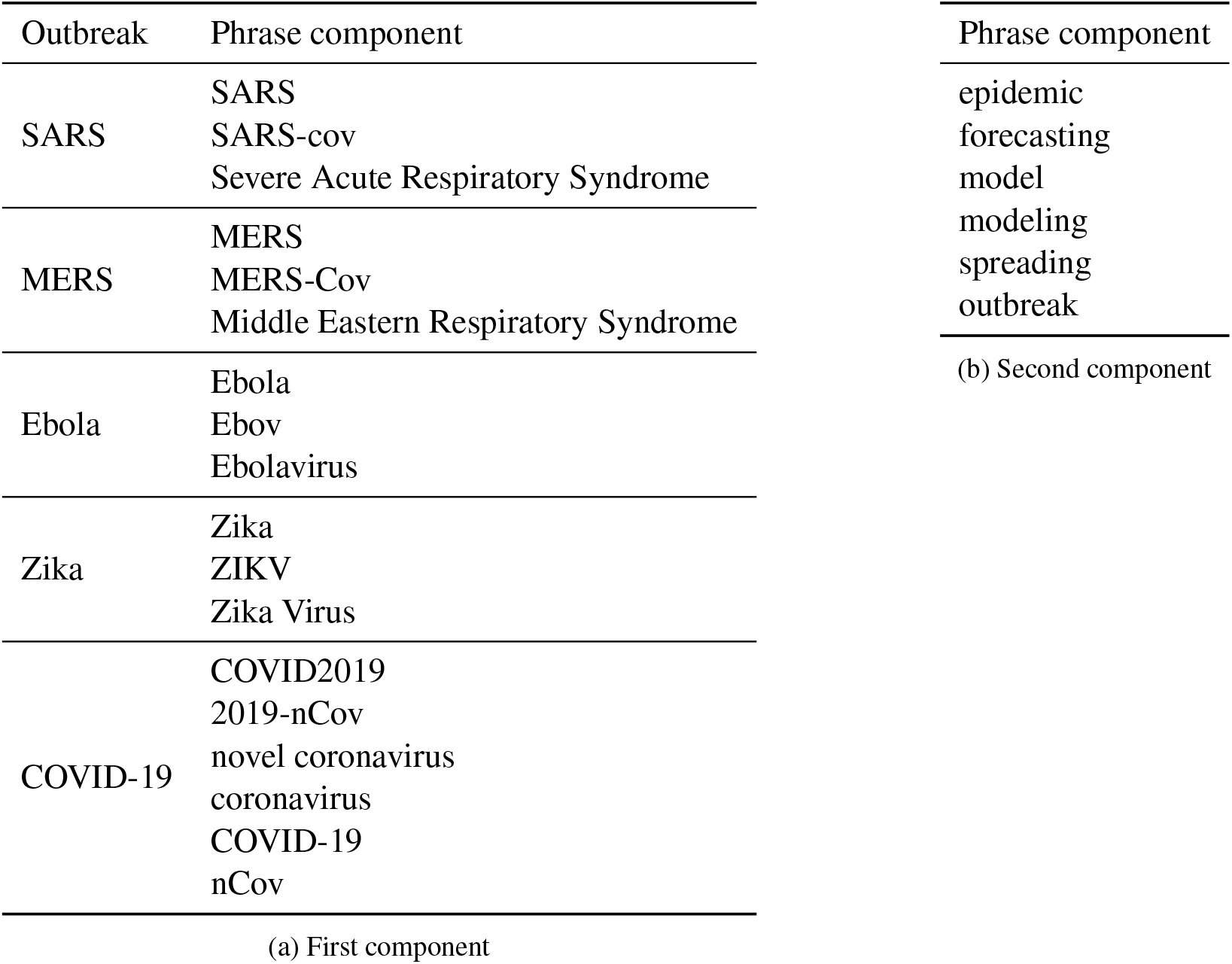
A reference table for constructing phrases used in 5.4. For each virus outbreak, the list of phrases consists of all combinations of the first component concatenated with the second component. For instance, ‘SARS epidemic’ or ‘COVID-19 forecasting’.

## Footnotes

1 https://github.com/seasonyao/BiorXivImpact

2 A minute percentage of titles are missing from the PubMed data. This is attributed to noisy data entry rather than some data being closed or open.

3 For computing time series, we simply count if a phrase occurred in a document independent of its importance score from TextRank. This approach scales easily to large corpora such as full text from PubMed papers.

4 A simple heuristic to get one changepoint is to use the first one, although this does not work universally well. Constraining WCPD to return a single changepoint (e.g. highest discrepancy) similarly does not consistently return the most desirable changepoints.

## References

J. Alexander, K. Bache, J. Chase, C. Freyman, J. D. Roessner, and P. Smyth. 2013. An exploratory study of interdisciplinarity and breakthrough ideas. In 2013 Proceedings of PICMET ’13: Technology Management in the IT-Driven Services (PICMET).

Jeffrey Alexander. 2013. A reasoning-based framework for the computation of technical emergence. GTM 2013-Atlanta, GA.

Samaneh Aminikhanghahi and Diane J. Cook. 2016. A survey of methods for time series change point detection. Knowledge and Information Systems, 51(2):339–367.

Jushan Bai. 1997. Estimating multiple breaks one at a time. Econometric Theory, 13(3):315–352.

Jeremy M. Berg, Needhi Bhalla, Philip E. Bourne, Martin Chalfie, David G. Drubin, James S. Fraser, Carol W. Greider, Michael Hendricks, Chonnettia Jones, Robert Kiley, Susan King, Marc W. Kirschner, Harlan M. Krumholz, Ruth Lehmann, Maria Leptin, Bernd Pulverer, Brooke Rosenzweig, John E. Spiro, Michael Stebbins, Carly Strasser, Sowmya Swaminathan, Paul Turner, Ronald D. Vale, K. VijayRaghavan, and Cynthia Wolberger. 2016. Preprints for the life sciences. Science, 352(6288).

Peter van den Besselaar and Ulf Sandström. 2018. Measuring researcher independence using bibliometric data: A proposal for a new performance indicator. Technical report, Cold Spring Harbor Laboratory.

Lutz Bornmann, K. Brad Wray, and Robin Haunschild. 2019. Citation concept analysis (CCA): a new form of citation analysis revealing the usefulness of concepts for other researchers illustrated by exemplary case studies including classic books by thomas s. kuhn and karl r. popper. Scientometrics, 122(2):1051–1074.

Charles LA Clarke, Nick Craswell, and Ian Soboroff. 2004. Overview of the TREC 2004 terabyte track. In TREC, volume 4.

Aaron Clauset, Daniel B. Larremore, and Roberta Sinatra. 2017. Data-driven predictions in the science of science. Science, 355(6324):477–480.

Susan E. Cozzens, Sonia Gatchair, Jongseok Kang, Kyung-Sup Kim, Hyuck Jai Lee, Gonzalo R. Ordóñez, and Alan L. Porter. 2010. Emerging technologies: quantitative identification and measurement. Techn. Analysis & Strat. Manag., 22.

Tirthankar Dasgupta and Lipika Dey. 2016. Automatic scoring for innovativeness of textual ideas. In Workshops at the Thirtieth AAAI Conference on Artificial Intelligence.

Philippe Desjardins-Proulx, Ethan P. White, Joel J. Adamson, Karthik Ram, Timotheée Poisot, and Dominique Gravel. 2013. The case for open preprints in biology. PLOS Biology, 11(5).

Laura Dietz, Steffen Bickel, and Tobias Scheffer. 2007. Unsupervised prediction of citation influences. In Proceedings of the 24th international conference on Machine learning - ICML’07. ACM Press.

Y. Dong, H. Ma, J. Tang, and K. Wang. 2018. Collaboration Diversity and Scientific Impact. ArXiv eprints.

Michael B. Eisen and Robert Tibshirani. 2020. How to identify flawed research before it becomes dangerous. New York Times. July 20.

Sergey Feldman, Kyle Lo, and Waleed Ammar. 2018. Citation count analysis for papers with preprints. ArXiv, abs/1805.05238.

Piotr Fryzlewicz. 2007. Unbalanced haar technique for nonparametric function estimation. Journal of the American Statistical Association, 102(480):1318–1327.

Eugene Garfield. 1967. Primordial concepts, citation indexing, and historio-bibliography. The Journal of library history, 2(3):235–249.

Eugene Garfield, A. I. Pudovkin, and V. S. Istomin. 2002. Algorithmic citation-linked historiography—–mapping the literature of science. Proceedings of the American Society for Information Science and Technology, 39(1):14–24.

Donna Harman. 2002. Overview of the TREC 2002 novelty track. In Proceedings of the Eleventh Text Retrieval Conference (TREC 2002), NIST Special Publication 500-251. Citeseer.

Drahomira Herrmannova, Petr Knoth, and Robert M. Patton. 2018a. Analyzing citation-distance networks for evaluating publication impact. In LREC.

Drahomira Herrmannova, Petr Knoth, Christopher Stahl, Robert Patton, and Jack Wells. 2018b. Text and graph based approach for analyzing patterns of research collaboration: An analysis of the TrueImpactDataset. In Proceedings of the Eleventh International Conference on Language Resources and Evaluation (LREC 2018), Paris, France. European Language Resources Association (ELRA).

B. Ian Hutchins, Xin Yuan, James M. Anderson, and George M. Santangelo. 2016. Relative citation ratio (RCR): A new metric that uses citation rates to measure influence at the article level. PLOS Biology, 14(9).

Iacopo Iacopini, Stasša Milojevicć, and Vito Latora. 2018. Network dynamics of innovation processes. Phys. Rev. Lett., 120.

David Jurgens, Srijan Kumar, Raine Hoover, Dan McFarland, and Dan Jurafsky. 2018. Measuring the evolution of a scientific field through citation frames. Transactions of the Association for Computational Linguistics, 6:391–406.

Margarita Karkali, François Rousseau, Alexandros Ntoulas, and Michalis Vazirgiannis. 2013. Efficient online novelty detection in news streams. In Lecture Notes in Computer Science, pages 57–71. Springer Berlin Heidelberg.

R. Killick, P. Fearnhead, and I. A. Eckley. 2012. Optimal detection of changepoints with a linear computational cost. Journal of the American Statistical Association, 107(500):1590–1598.

Daniel King, Doug Downey, and Daniel S. Weld. 2020. High-Precision Extraction of Emerging Concepts from Sientific Literature. In Proceedings of the 43rd International ACM SIGIR Conference on Research and Development in Information Retrieval (SIGIR’20), Virtual Event, China. ACM.

Richard Klavans, Kevin W. Boyack, and Dewey A. Murdick. 2020. A novel approach to predicting exceptional growth in research. arXiv e-prints, page arXiv:2004.13159.

Peter Klimek, Aleksandar S. Jovanovic, Rainer Egloff, and Reto Schneider. 2016. Successful fish go with the flow: citation impact prediction based on centrality measures for term–document networks. Scientometrics, 107(3):1265–1282.

Harlan M. Krumholz, Theodora Bloom, and Joseph S. Ross. 2020. Preprints can fill a void in times of rapidly changing science. StatNews.

Bruno Latour and Steve Woolgar. 1986. Laboratory Life: The Construction of Scientific Facts. Princeton University Press, Princeton, NJ.

Michael S Lauer, Harlan M Krumholz, and Eric J Topol. 2015. Time for a prepublication culture in clinical research? Lancet, 386:2447–2449.

Cynthia Lokker, K Ann McKibbon, R James McKinlay, Nancy L Wilczynski, and R Brian Haynes. 2008. Prediction of citation counts for clinical articles at two years using data available within three weeks of publication: retrospective cohort study. BMJ, 336(7645):655–657.

Kathy McKeown, Hal Daume, Snigdha Chaturvedi, John Paparrizos, Kapil Thadani, Pablo Barrio, Or Biran, Suvarna Bothe, Michael Collins, Kenneth R. Fleischmann, Luis Gravano, Rahul Jha, Ben King, Kevin McInerney, Taesun Moon, Arvind Neelakantan, Diarmuid O’Seaghdha, Dragomir Radev, Clay Templeton, and Simone Teufel. 2016. Predicting the impact of scientific concepts using full-text features. Journal of the Association for Information Science and Technology, 67(11):2684–2696.

Rada Mihalcea and Paul Tarau. 2004. Textrank: Bringing order into text. In Proceedings of the 2004 conference on empirical methods in natural language processing.

Paco Nathan. 2016. Pytextrank, a python implementation of textrank for phrase extraction and summarization of text documents. https://github.com/DerwenAI/pytextrank/.

Larry Peiperl. 2018. Preprints in medical research: Progress and principles. PLOS Medicine, 15(4).

Brandon K. Peoples, Stephen R. Midway, Dana Sackett, Abigail Lynch, and Patrick B. Cooney. 2017. Twitter predicts citation rates of ecological research. PLoS ONE, 11.

Kendall Powell. 2016. Does it take too long to publish research? Nature, 530.

Lindor Qunaj, Raina H. Jain, Coral L. Atoria, Renee L. Gennarelli, Jennifer E. Miller, and Peter B. Bach. 2018. Delays in the publication of important clinical trial findings in oncology. JAMA Oncology, 4(7).

Daniele Rotolo, Diana Hicks, and Ben Martin. 2015. What is an emerging technology? SSRN Electronic Journal.

Andrey Rzhetsky, Jacob G. Foster, Ian T. Foster, and James A. Evans. 2015. Choosing experiments to accelerate collective discovery. Proceedings of the National Academy of Sciences, 112(47):14569–14574.

Angelo A. Salatino, Francesco Osborne, and Enrico Motta. 2018. AUGUR. In Proceedings of the 18th ACM/IEEE on Joint Conference on Digital Libraries. ACM.

Sarvenaz Sarabipour, Humberto J. Debat, Edward Emmott, Steven J. Burgess, Benjamin Schwessinger, and Zach Hensel. 2019. On the value of preprints: An early career researcher perspective. PLOS Biology, 17(2).

Patrick D. Schloss. 2017. Preprinting microbiology. mBio, 8(3).

András P. Schubert and Gábor A. Schubert. 1997. Inorganica Chimica Acta: its publications, references and citations. an update for 1995–1996. Inorganica Chimica Acta, 266(2):125 – 133.

Dafna Shahaf, Carlos Guestrin, and Eric Horvitz. 2012. Metro maps of science. In Proceedings of the 18th ACM SIGKDD International Conference on Knowledge Discovery and Data Mining, KDD ’12, New York, NY, USA. ACM.

Xiaolin Shi, Jure Leskovec, and Daniel A. McFarland. 2010. Citing for high impact. CoRR, abs/1004.3351.

Sotaro Shibayama and Jian Wang. 2020. Measuring originality in science. Scientometrics, 122.

R. Sinatra, D. Wang, P. Deville, C. Song, and A.-L. Barabasi. 2016. Quantifying the evolu tion of individual scientific impact. Science, 354(6312):aaf5239–aaf5239.

Henry Small, Kevin W. Boyack, and Richard Klavans. 2019. Citations and certainty: a new interpretation of citation counts. Scientometrics, 118(3):1079–1092.

Ian Soboroff and Donna Harman. 2003. Overview of the TREC 2003 novelty track. In TREC. Citeseer.

Ian Soboroff and Donna Harman. 2005. Novelty detection: the TREC experience. In Proceedings of the conference on Human Language Technology and Empirical Methods in Natural Language Processing. Association for Computational Linguistics.

Iman Tahamtan, Askar Safipour Afshar, and Khadijeh Ahamdzadeh. 2016. Factors affecting number of citations: a comprehensive review of the literature. Scientometrics, 107(3):1195–1225.

Derek Tam, Nicholas Monath, Ari Kobren, Aaron Traylor, Rajarshi Das, and Andrew McCallum. 2019. Optimal transport-based alignment of learned character representations for string similarity. In Proceedings of the 57th Annual Meeting of the Association for Computational Linguistics, pages 5907–5917, Florence, Italy. Association for Computational Linguistics.

Charles Truong, Laurent Oudre, and Nicolas Vayatis. 2020. Selective review of offline change point detection methods. Signal Processing, 167:107299.

Brian Uzzi, Satyam Mukherjee, Michael Stringer, and Ben Jones. 2013. Atypical combinations and scientific impact. Science, 342(6157):468–472.

Ronald D. Vale. 2015. Accelerating scientific publication in biology. Proceedings of the National Academy of Sciences, 112(44).

Arnout Verheij, Allard Kleijn, Flavius Frasincar, and Frederik Hogenboom. 2012. A comparison study for novelty control mechanisms applied to web news stories. In 2012 IEEE/WIC/ACM International Conferences on Web Intelligence and Intelligent Agent Technology. IEEE.

Jian Wang, Reinhilde Veugelers, and Paula Stephan. 2016. Bias against novelty in science: A cautionary tale for users of bibliometric indicators. Technical report, National Bureau of Economic Research.

Lucy Lu Wang, Kyle Lo, Yoganand Chandrasekhar, Russell Reas, Jiangjiang Yang, Darrin Eide, Kathryn Funk, Rodney Kinney, Ziyang Liu, William Merrill, Paul Mooney, Dewey Murdick, Devvret Rishi, Jerry Sheehan, Zhihong Shen, Brandon Stilson, Alex D. Wade, Kuansan Wang, Chris Wilhelm, Boya Xie, Douglas Raymond, Daniel S. Weld, Oren Etzioni, and Sebastian Kohlmeier. 2020. CORD-19: The Covid-19 open research dataset. arXiv e-prints, page arXiv:2004.10706.

Mengyang Wang and Lihe Chai. 2018. Three new bibliometric indicators/approaches derived from keyword analysis. Technical Report 2, Springer Science and Business Media LLC.

Ian Wesley-Smith, Carl T. Bergstrom, and Jevin D. West. 2016. Static ranking of scholarly papers using article-level eigenfactor (alef). ArXiv, abs/1606.08534.

Fen Zhao, Yi Zhang, Jianguo Lu, and Ofer Shai. 2019. Measuring academic influence using heterogeneous author-citation networks. Scientometrics, 118(3):1119–1140.

